# ModrRNA1: A tool for possible RNA modification site identification with associated putative enzymes in bacterial rRNA

**DOI:** 10.1101/2023.11.01.564906

**Authors:** Uvindu Thilanka, Paaramitha Sithumini Warushavithana

**Affiliations:** BioTechie; University of Colombo

**Keywords:** ModrRNA1, Bacterial rRNA, rRNA modifications, rRNA modification enzymes, modification site annotation, methyltransferases, structural comparisons, RNA methylation

## Abstract

Ribosomal RNA modifications play a crucial role in bacteria, impacting function, ribosome formation, and antibiotic resistance. ModrRNA1 replaces manual methods and introduces a user-friendly, streamlined bioinformatics tool for the identification of potential rRNA modification sites and associated enzymes, across various bacteria.

This tool utilizes a curated database of known bacterial rRNA modifications and enzymes, employing sequence alignment with a customized scoring system for precise site identification. High-confidence alignments generate annotated RNA sequence plots. Subsequently, enzyme detection is facilitated through structural comparisons, phylogenetics, and BLAST.

ModrRNA1 boasts a user-friendly web interface hosted at https://modrrna.biotechie.org with the open-source code for customization available at https://github.com/sciguysl/ModrRNA1. The results demonstrate ModrRNA1’s efficacy in identifying potential non-species-specific bacterial rRNA modification sites and species-specific associated enzymes.

## Introduction

Nucleotide modifications of ribosomal RNA (rRNA) can occur post-transcriptionally through three main pathways—namely, conversion of uridine to pseudouridine (5-ribosyl uracil), methylation of 2’ hydroxyls, and base alteration; most of which are followed by methylation at various positions [1]. The distribution of all *Escherichia coli* rRNA modifications in the ribosome has been identified and localized [1]. Modifications mostly occur in the interior of the RNA mass of the ribosomes, oriented towards the faces of the ribosomal subunits, and they are distributed mostly in conserved regions of both the large and small subunits. Hence, it is apparent that modifications affect not only the structure but also the ribosomal functions [1]. In contrast, since modified nucleotides are absent from regions where ribosomal proteins are abundant, modifications do not directly impact RNA-protein interactions.

rRNA modifications commonly seen in *E. coli* are carried out by site-specific or region-specific protein-only enzymes. Almost all rRNA modifications of the 70S ribosome of *E. coli* are known and there are 36 modified nucleotides in total [2]. The number of methylated and pseudouridylated nucleotides found in 16S rRNA of *E. coli* are 10 and 1, respectively. Modified nucleotides of 23S rRNA in *E. coli* are 13 methylated ones, 9 pseudouridines, 1 methyl pseudouridine, 1 dihydrouridine, and 1 hydroxycytidine. Positions of modified nucleotides in the *E. coli* 16S and 23S rRNA and the enzymes responsible for their formation are shown below. (Tables 1 and 2) [3].

**Table 1:**
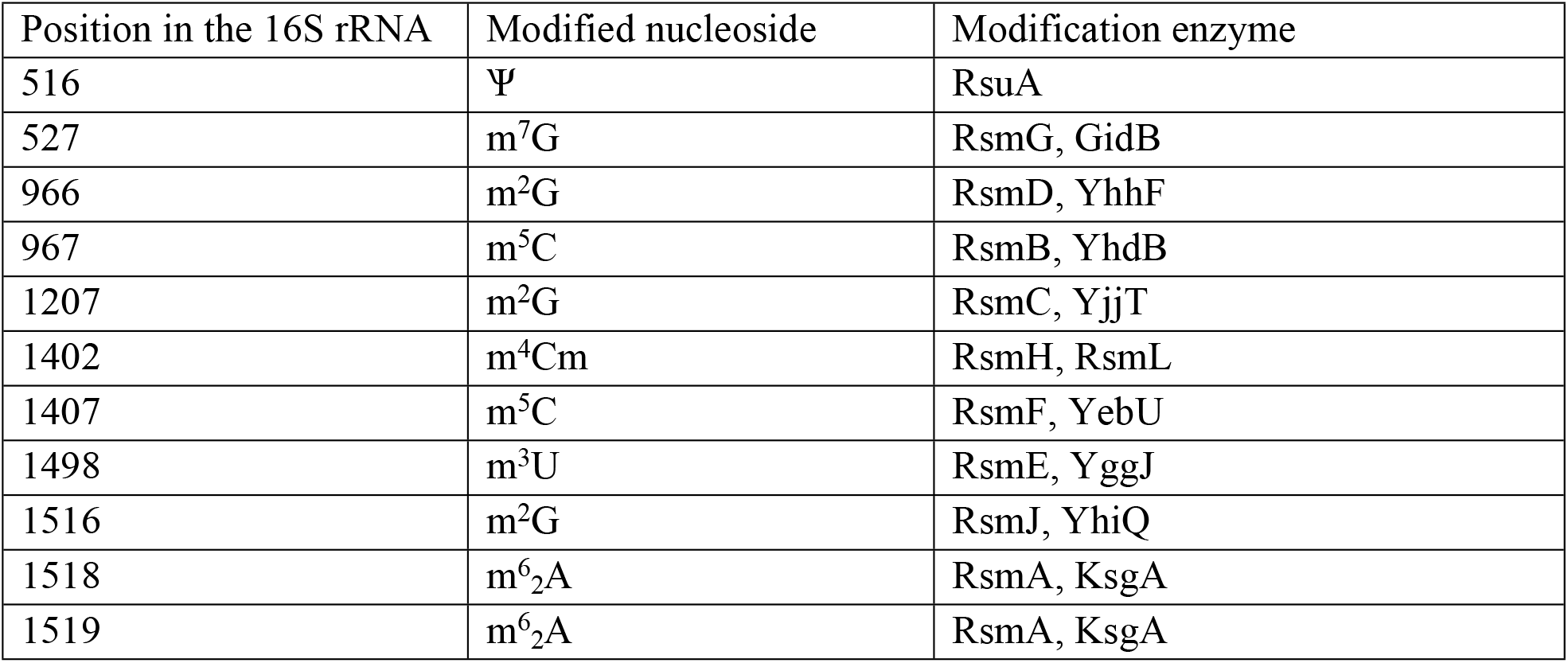
Modified nucleotides of 16S rRNA of E. coli and associated modifying enzymes.

| Position in the 16S rRNA | Modified nucleoside | Modification enzyme |
| --- | --- | --- |
| 516 | Ψ | RsuA |
| 527 | m<sup>7</sup>G | RsmG, GidB |
| 966 | m<sup>2</sup>G | RsmD, YhhF |
| 967 | m<sup>5</sup>C | RsmB, YhdB |
| 1207 | m<sup>2</sup>G | RsmC, YjjT |
| 1402 | m<sup>4</sup>Cm | RsmH, RsmL |
| 1407 | m<sup>5</sup>C | RsmF, YebU |
| 1498 | m<sup>3</sup>U | RsmE, YggJ |
| 1516 | m<sup>2</sup>G | RsmJ, YhiQ |
| 1518 | m<sup>6</sup><sub>2</sub>A | RsmA, KsgA |
| 1519 | m<sup>6</sup><sub>2</sub>A | RsmA, KsgA |

**Table 2:** Modified nucleotides of 23S rRNA of E. coli and associated modifying enzymes.

| Position in the 23S rRNA | Modified nucleoside | Modification enzyme |
| --- | --- | --- |
| 745 | m<sup>1</sup>G | RrmA, YebH |
| 746 | Ψ | RluA |
| 747 | m<sup>5</sup>U | YbjF, RumB |
| 955 | Ψ | RluC |
| 1618 | m<sup>6</sup>A | YbiN |
| 1835 | m<sup>2</sup>G | RlmG, YgjO |
| 1911 | Ψ | RluD |
| 1915 | m<sup>3</sup>Ψ | RluD, YbeA |
| 1917 | Ψ | RluD |
| 1939 | m<sup>5</sup>U | RlmD |
| 1962 | m<sup>5</sup>C | RlmL |
| 2030 | m<sup>6</sup>A | YhiR, RlmJ |
| 2069 | m<sup>7</sup>G | RlmKL |
| 2251 | Gm | RlmB |
| 2445 | m<sup>2</sup>G | RlmL |
| 2449 | hU |  |
| 2457 | Ψ | RluE |
| 2498 | Cm | RlmM |
| 2501 | oh<sup>5</sup>C |  |
| 2503 | m<sup>2</sup>A | RlmN |
| 2504 | Ψ | RluC |
| 2552 | Um | RlmE, RrmJ |
| 2580 | Ψ | RluC |
| 2604 | Ψ | RluF |
| 2605 | Ψ | RluB |

In bacteria, the most prominent type of nucleotide modification is methylation. Additionally, most methylated nucleotides are found exposed on the surface of ligand-free small subunits to be readily accessible to methyltransferases. Some enzymes need ribosomal proteins to carry out modifications while other enzymes only modify while rRNA is protein free. In vitro, sub-ribosomal particle analysis has shown that most methylations occur during the early or late steps of ribosome assembly [4]. Various Pseudouridine synthases carry out pseudouridylation.

Pseudouridylation leads to increased structural rigidity of the sugar-phosphate backbone, alteration of base stacking interactions [5], steric properties, and different H-bonding patterns [1]. Both pseudouridylation and methylation of 2’ hydroxyls cause enhanced structural rigidity for single-stranded and double-stranded regions [1]. Methylation of 2′-OH prevents hydrolysis of inter-nucleotide bonds and could influence base stacking [3]. Such effects on the chemical properties in turn help RNA folding by resolving misfolded structures or by preventing their formation and change the assembly and activity of the ribosome.

Methyltransferase-mediated rRNA modifications play a vital role in the generation of functional ribosomes. A unique modified cytosine found in almost all bacterial 16S rRNA which is methylated at the m^4^Cm1402, is important for P-site structure formation and enhances the accuracy of initiation codon recognition [6]. Another highly conserved modification is the dimethylation of two adenosines carried out by Methyltransferase KsgA. X-ray crystal structure of the *Thermus thermophilus* 30S ribosomal subunit shows that defective KsgA methyltransferase results in the absence of methylation, which causes translational mistakes in the initiation and elongation stages of protein synthesis and confers resistance to the antibiotic kasugamycin, due to a conformation change that affect the antibiotic binding [7]. Moreover, mutations in the gene that codes for methyltransferase RsmG lead to streptomycin resistance [8]. Therefore, it is likely that the mechanisms of a significant number of antibiotic resistances arise from rRNA methylations at specific nucleotides.

Deletions of *rsmB* and *rsmD* genes have been shown to cause disruption of transcription, and misregulation of attenuation control of tryptophan operon in *E. coli* [9]. Mutations in methyltransferase genes have also been proven to cause moderate to drastic cell growth retardation respectively in the cases of methyltransferase RlmF [10] and RrmJ [11]. Methyltransferase orthologs may possess functions in addition to their rRNA methyltransferase activity such as acting as RNA chaperones, as seen in KsgA. This enzyme also plays a role in the suppression of coldsensitivity and may be involved in the acid shock response of *E. coli* [12].

In summary, these rRNA modifications play a pivotal role in bacteria. Therefore, A tool for general identification of modification sites and the enzymes associated with each modification can be important in studying the formation and functionality of a ribosome of a given bacteria species, and in gaining clinically relevant insight into what confers antibiotic resistance in bacteria by better understanding rRNA modifications. The precise localization of these sites and the identification of potential enzymes are essential for deciphering their biological significance. Existing tools for this purpose are lacking, leading researchers to rely on manual methods. ModrRNA1 integrates tasks from the general identification of potential rRNA modification sites to shortlisting putative enzymes and performing BLAST analysis [13] on the query bacterial genome, thereby streamlining and simplifying the process for researchers in a user-friendly manner. This tool can be utilized in finding uncharacterized probable rRNA modification sites and previously unacknowledged RNA modification enzymes in a variety of bacteria and to find proteins that have RNA modification functionality.

## Method

### Database Preparation

ModrRNA1 is an open-source web application developed in Python 3. It utilizes a database of bacterial rRNA modifications, which consists of conserved sequences located both downstream and upstream of each modification site, along with the details on the respective modification enzymes. This database was curated by collating data from the *RMBase* [14] bacterial RNA modification site datasets and cross-referencing it with information from *MODOMICS* [15] for annotation with the associated enzymes. In addition, a protein structure library was created using *ESMFold* [16] for all the enzymes involved in these reference modifications. (Figure 1)

**Figure 1:**
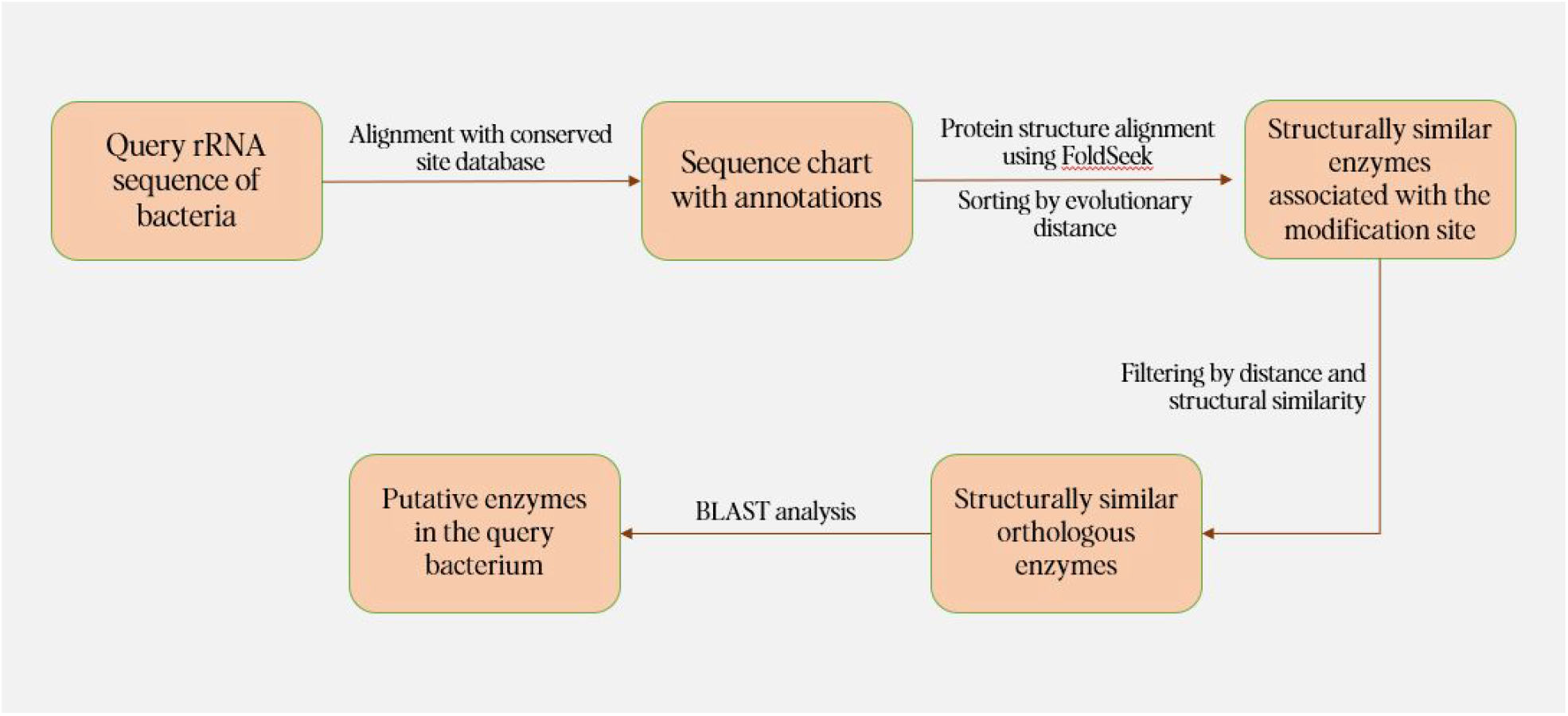
A flowchart depicting the workflow of ModrRNA1.

### Sequence Alignment

In ModrRNA1, firstly the modification site sequences are duplicated and transformed to their reverse complements using the *Bio*.*Seq* module in *Biopython* [17]. Then it aligns these sequences to the input query sequence using the *Bio*.*pairwise2* module. We customized the scoring function of the aligner to give 2 points for an identical character, -1 point for a non-identical character, -4 points for a gap opening, and -1 point for a gap extension to minimize gap formation when aligning. An accuracy score is calculated as a percentage to assess the quality of the alignment using the simple equation below.

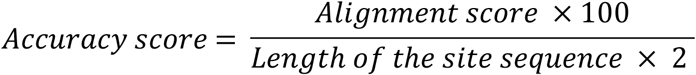

### Site Annotation

The tool removes redundant sequences and obtains alignments of high confidence by applying a threshold of an accuracy score greater than or equal to 60. Based on this data, it generates a single-axis plot with the query sequence on the x-axis using the *DnaFeaturesViewer* library [18]. The detected RNA modifications are annotated visually and are distinguishable at their respective sites on the plot. These annotations include the type of modification, nucleotide location, and the unique identifier used to find the enzyme(s) in subsequent steps. The *Bio*.*SeqIO* package, in conjunction with the *Bio*.*SeqFeature* module is employed to tag these annotations and output them into GenBank and CSV format, which can be downloaded by the user and utilized in external analyses.

### Enzyme Detection

If the source bacterium of the query sequence is known, ModrRNA1 allows the user to find the most likely enzymes involved in a particular modification. The specific enzyme structures are screened using the *Protein Data Bank (PDB)* [19] and virtual protein structure databases (*AlphaFold*) [20] to screen structurally similar enzymes, employing the *FoldSeek* API [21]. Thresholds greater than or equal to 50% of query coverage (50 ≤ qscore) and 60 ≤ TM-score normalized by the query length (prob) are applied to filter and shortlist the results.

The evolutionary distance is calculated based on the topology (branching structure of the tree) between the query bacterium and the source bacteria of these enzymes using the *NCBITaxa* module in the *ete3* library [22]. The most phylogenetically related enzymes are selected and sorted by comparing the evolutionary distance. The results are outputted as entries to the Protein Data Bank and the AlphaFold Protein Structure Database (AFDB) [20]. The user can compare these calculated distances, query coverage scores, and TM-scores (prob) to locate probable enzymes and then perform a blastp or a tblastn analysis [13] within this tool itself on the query bacterial genome. In cases where high-scoring alignments are found in the BLAST search, it suggests the presence of the specific rRNA modification-associated enzyme and the possible modification of the respective site in the queried bacteria. (Figure 1)

### Web Application

*Streamlit*, which is an open-source Python-based application framework was used to create the web application. A user-friendly UI with clear instructions was implemented, ensuring accessibility to a broad user base. It is hosted in an Azure app service instance in Linux with Python 3.8 version.

### Testing

The tool was tested to assess its operational functionality, employing 16S ribosomal RNA (rRNA) sequences derived from *Escherichia sp*. strain Esraa 4 (MT647245.1) and *Thermus thermophilus* HB8 (NR_037066.1). The potential modification sites identified in ModrRNA1 were compared with known rRNA modification sites obtained from MODOMICS. As the positions are formatted in *E. coli* numbering, the *T. thermophilus* 16S rRNA annotated Genbank file obtained from ModrRNA1 was aligned with the *E. coli* 16S rRNA sequence using Benchling to locate the positions.

For tool evaluation, it was applied to 23S rRNA sequences derived from organisms significantly divergent from the source bacteria from which the initial conserved site sequences were obtained. The query sources were *Deinococcus radiodurans* strain R1 (NR_076078.2) and *Haloarcula marismortui* ATCC 43049 (NR_076318.1). Since the positions are *E. coli*-numbered, the query sequences were aligned with the 23S rRNA of *E. coli*. The results were tabulated in Table 4. To evaluate the Enzyme Explorer in ModrRNA1, six randomly selected potential modification sites in *T. Thermophilus* were processed, specifically: ψ (499), m^2^_2_ G (943), m^5^C (1378), m^6^_2_ A (1491), m^4^Cm (1380), and m^3^U (1471). A blastp search was done using the top match which was selected out of FoldSeek shortlisted hits based on evolutionary distance. The BLAST alignment yielding the highest score was postulated as the putative enzyme, and the results for each site were tabulated in Table 5.

**Table 3:** Known 16S rRNA modification sites and ModrRNA1 fetched sites comparison between E. coli and T. Thermophilus. (Modification positions in the first column are in E. coli numbering)

| Modification position | Modification (known) in *E. coli* | Modification (ModrRNA1) in *E. coli* | Modification (known) in *T. Thermophilus* | Modification (ModrRNA1) in *T. Thermophilus* |
| --- | --- | --- | --- | --- |
| 516 | Ψ | Ψ | Ψ | Ψ (499) |
| 527 | m<sup>7</sup>G | m<sup>7</sup>G | m<sup>7</sup>G | m<sup>7</sup>G (510) |
| 966 | m<sup>2</sup>G | m<sup>2</sup>G | m<sup>2</sup><sub>2</sub>G | m<sup>2</sup><sub>2</sub>G (943) |
| 967 | m<sup>5</sup>C | m<sup>5</sup>C | m<sup>5</sup>C | m<sup>5</sup>C (944) |
| 1207 | m<sup>2</sup>G | m<sup>2</sup>G | m<sup>2</sup>G | m<sup>2</sup>G (1186) |
| 1400 | - | - | m<sup>5</sup>C | m<sup>5</sup>C (1378) |
| 1402 | m<sup>4</sup>Cm | m<sup>4</sup>Cm | m<sup>4</sup>Cm | m<sup>4</sup>Cm (1380) |
| 1404 | - | - | m<sup>5</sup>C | m<sup>5</sup>C (1382) |
| 1407 | m<sup>5</sup>C | m<sup>5</sup>C | m<sup>5</sup>C | m<sup>5</sup>C (1385) |
| 1498 | m<sup>3</sup>U | m<sup>3</sup>U | m<sup>3</sup>U | m<sup>3</sup>U (1471) |
| 1516 | m<sup>2</sup>G | m<sup>2</sup>G | - | - |
| 1518 | m<sup>6</sup><sub>2</sub>A | m<sup>6</sup><sub>2</sub>A | m<sup>6</sup><sub>2</sub>A | m<sup>6</sup><sub>2</sub>A (1491) |
| 1519 | m<sup>6</sup><sub>2</sub>A | m<sup>6</sup><sub>2</sub>A | m<sup>6</sup><sub>2</sub>A | m<sup>6</sup><sub>2</sub>A (1492) |
| 1540 | - | - | Ψ | Ψ (1513) |
| 1541 | - | - | Ψ | Ψ (1514) |

**Table 4:**
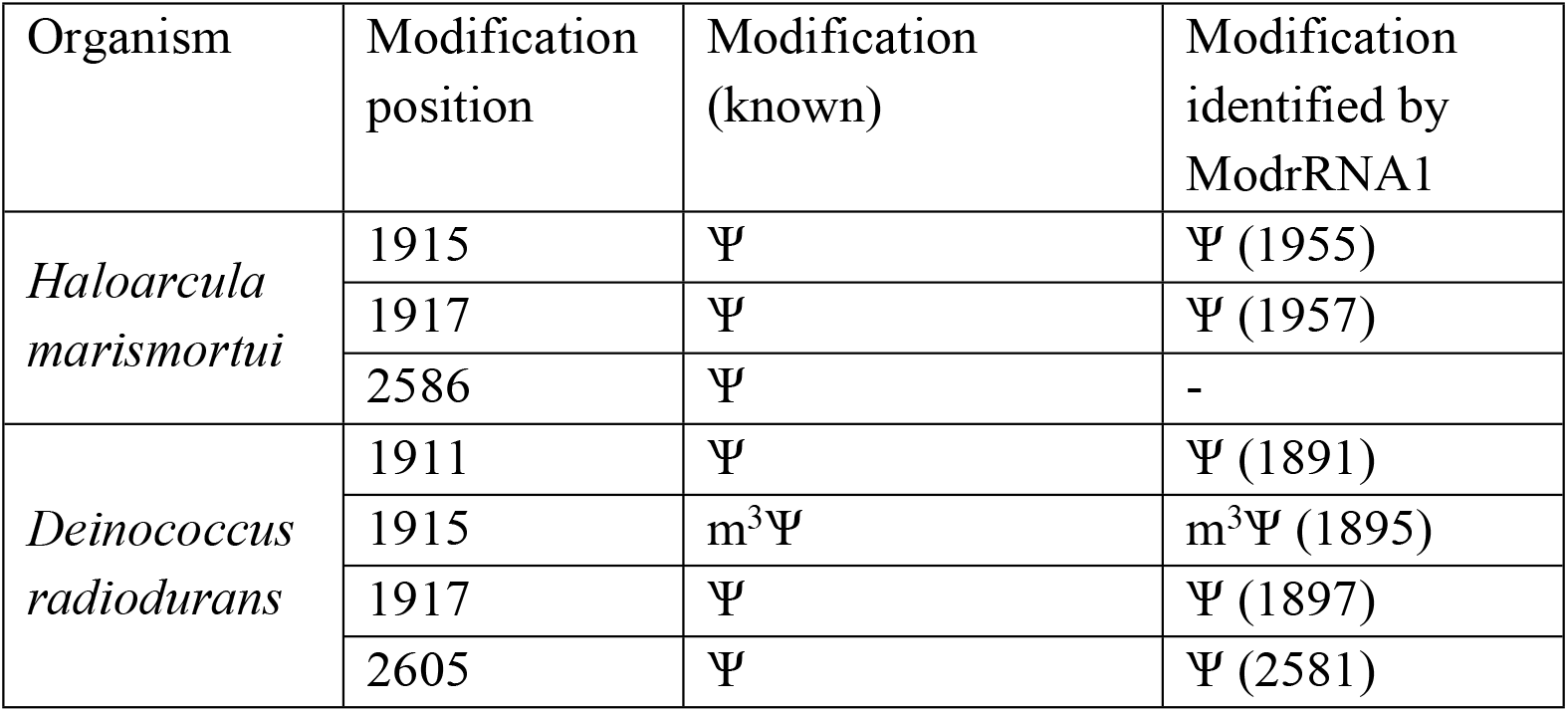
Known 23S rRNA modification sites and ModrRNA1 fetched sites comparison. (Modification positions in the first column are in E. coli numbering.)

**Table 5:**
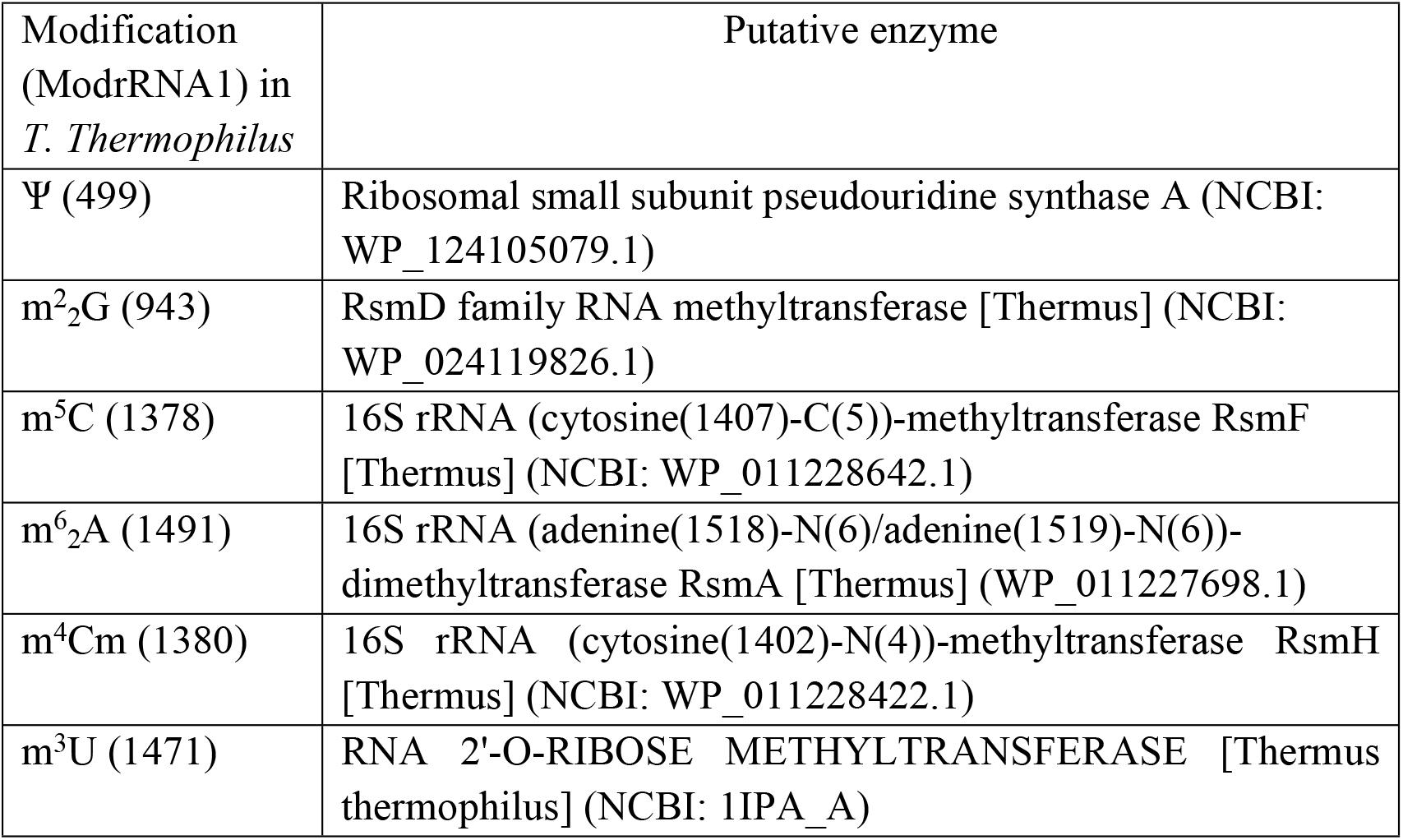
ModrRNA1 BLAST results of several modification sites in T. Thermophilus 16S rRNA.

## Results and discussion

As shown in the comparison in Table 3, all the characterized 16S rRNA modification sites of *T. thermophilus* and *E. coli* are identified by ModrRNA1 as intended. This encompasses the identification of several types of modifications including methylations of different nucleotides (m^2^G, m^7^G, m^4^C, m^5^C, m^3^U, m^6^_2_A) and also pseudouridinylations, in both bacterial species. There were several extra potential modification sites identified by the tool as in this step the code aligns all the conserved sequences from the site database. These sites may or may not represent actual modification sites, and their confirmation may require subsequent enzyme search steps. In the evaluation results presented in Table 4, we compared the modifications detected by the tool with those reported in a previous study. All pseudouridylation sites were successfully identified, with the exception of the one located at position 2586 (*E. coli* numbering). This discrepancy might be attributed to variations in the query sequence of *Haloarcula marismortui* used during the evaluation, as the exact 23S rRNA sequence employed had not been disclosed in this study. It should be noted that the absence of certain missing sites may be due to their non-inclusion of the specific conserved site sequences in the ModrRNA database. This database exclusively comprises sequences that have been confirmed through multiple studies. Therefore, if a new modification site is identified in a single study, it may not necessarily appear as a potential modification site in this tool. To address this limitation, our objective is to maintain an up-to-date database by continuously incorporating newly confirmed (through more than one study) site sequences. In the enzyme evaluation presented in Table 5, it is clear that the putative enzymes identified exhibit a consistent correlation with the associated modification types, as evidenced by the comparisons with Table 1 and Table 2 and further literature. The methyltransferase RsmF of *T. thermophilus* has been identified by multiple studies as responsible for multi-site-specific modifications: m^5^C1407 (1385), m^5^C1400 (1378), and m^5^C1404 (1382) [23,24]. Methyltransferase RsmH which is responsible for the N^4^-methylation of C1402 (1380) in *E. coli*[25] 16S rRNA and RsmH is also conserved in *T. Thermophilus* [26]. Dimethyltransferase RsmA (KsgA) carries out the most highly conserved rRNA modifications; A1518 (1491) and A1519 (1492) of 16S rRNA [27]. One study found the crystal structure of *Thermus thermophilus* hypothetical RNA 2’-O-ribose methyltransferase that seems corresponds with m^3^U (1471) modification [28].

The ability of ModrRNA1 to find potential modification sites and associated enzymes within bacterial rRNA sequences as demonstrated through the aforementioned results has wide-reaching implications. It can streamline the preliminary bioinformatics steps involved in rRNA modification enzyme research, providing a user-friendly platform for researchers to explore uncharacterized modification sites and enzymes across diverse bacteria, eliminating the need for manual methods that researchers usually rely upon. The open-source codebase, available on GitHub without restrictions, can be utilized to develop novel tools built upon this pipeline. These tools can be designed to target enzymes that operate on conserved sequences, similar to the methodology employed in this study.

## Conclusion

ModrRNA1 appears to be effective in identifying potential rRNA modification sites in a non-species-specific manner and associated enzymes specific to the query bacteria. It streamlines bioinformatics processes, offering a user-friendly alternative to manual methods. This tool may aid in research that advances our understanding of rRNA modifications and their functional implications in various biological contexts.

### Future perspective

Future directions involve expanding this tool to organisms beyond bacteria and incorporating other RNA types. Additionally, the database of conserved modification sequences is to be improved continuously.

Future studies will also focus on improving ModrRNA1 to incorporate experimental data, such as mass spectrometry or next-generation sequencing for the identification of putative rRNA modifications. The streamlined approach of the tool may also benefit from the incorporation of machine learning models for faster and more accurate prediction of modification sites based on sequence features.

### Summary points

- Ribosomal RNA modifications are pivotal to many bacterial functions.
- ModrRNA1 employs a database of known rRNA modification sites and the associated enzymes.
- The database is then utilized for sequence alignment, customized scoring, and generation of annotated plots over a given RNA sequence.
- Associated enzymes are detected through structural comparisons, phylogenetics and BLAST searches.
- Web application is user-friendly UI, open-source, Python-based, and hosted in an Azure app.
- The tool’s operational functionality was tested by various methods.
- ModrRNA1’s effectively identifies non-species-specific rRNA modification sites and associated enzymes in diverse bacteria.
- Provides researchers a user-friendly interface to discover new insights into bacterial ribosome functionality and RNA modification enzymes.

## Notes

https://modrrna.biotechie.org/

### Competing Interest Statement

The authors have declared no competing interest.

### Summary of Updates

The newest version that is undergoing peer-review in the BioTechniques journal is uploaded.

